# A Simple Approximation To The Bias Of Gene-Envinornment Interactions In Case/Control Studies With Silent Disease

**DOI:** 10.1101/444596

**Authors:** Iryna Lobach, Joshua Sampson, Siarhei Lobach, Alexander Alekseyenko, Alexandra Pryatinska, Tao He, Li Zhang

## Abstract

One of the most important research areas in case-control Genome-Wide Association Studies is to determine how the effect of a genotype varies across the environment or to measure the gene-environment interaction (GxE). We consider the scenario when some of the “healthy” controls actually have the disease and when the frequency of these latent cases varies by the environmental variable of interest. In this scenario, performing logistic regression of clinically defined case status on the genetic variant, environmental variable, and their interaction will result in biased estimates of GxE interaction. Here, we derive a general theoretical approximation to the bias in the estimates of the GxE interaction and show, through extensive simulation, that this approximation is accurate in finite samples. Moreover, we apply this approximation to evaluate the bias in the effect estimates of the genetic variants related to mitochondrial proteins a large-scale Prostate Cancer study.

## INTRODUCTION

One major objective in case-control Genome-Wide Association Studies (GWAS) is to determine how the effect of a genotype varies across the environment, i.e. to measure the gene-environment interaction (GxE). Understanding the GxE interaction can provide valuable clues into the underlying pathophysiologic mechanism of complex diseases (Ritz et al, 2017). A major complication is that supposedly “healthy” controls are often undiagnosed cases and the frequency of these latent cases may vary by environmental variables. Hence, the estimated GxE interaction with respect to the *true* pathophysiologic disease status would be biased if the analyses used only the clinically diagnosed disease status. The problem of latent cases is relatively common. For example, Atrial Fibrillation is undiagnosed in 5-17% of the population above the age of 75 (Panisello-Tafalla et al. 2015), non-alcoholic fatty liver disease is undiagnosed in 14-30% of the adult population (El-Kader et al., 2015), and acute coronary thrombosis is undiagnosed in >10% of individuals at the time of death (Anderson et al, 1989). Our specific motivating example is a large GWAS of prostate cancer. At autopsy, approximately 29%, 36%, and 47% of “healthy” men aged 60-69, 70-79 and 80+ years have undiagnosed prostate cancer, with the exact frequencies varying by race and ethnicity (Jahn et al, 2015).

We illustrate below why the GxE can appear to be associated with the disease status if presence of the silent cases is ignored based on a hypothetical example.

Shown on **Figure 1** is an example when frequency of a minor allele does differ by the true diagnosis defined as *D* = 0 to indicate controls, *D* = 1^*^ silent disease and *D* = 1 cases, but not by the environmental variable *X* = 1,2. But because diagnosed controls are in fact silent cases when *X* = 1, and 30% of the controls frequency of the silent disease varies by the environment (10% of clinically are silent cases when *X* = 2), there appears to be GxE on the clinical diagnosis.

**Figure 1:**
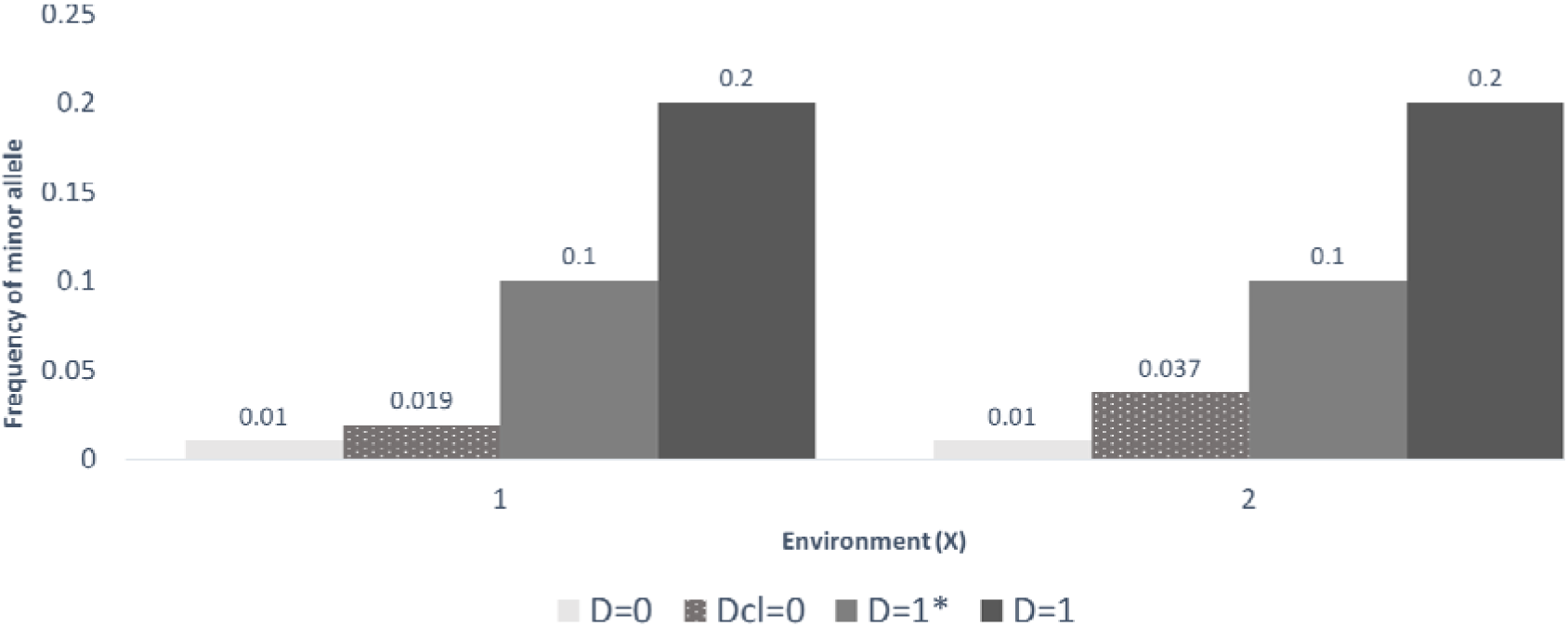
Frequency of the minor allele by a binary environmental variable ( ) on the x-axis for the *true* disease state (controls:, silent disease and case) and for the clinically diagnosed status that includes both *true* controls and silent cases. Shown is a hypothetical example when frequencies of the minor allele do not differ by the environmental variable on the *true* disease status and genotype is associated with the *true* disease status. Because frequency of the silent disease within the set of clinically diagnosed controls varies by the environment (10% of clinically diagnosed controls are in fact silent cases when, and 30% of the controls are silent cases when), there appears to be GxE on the clinical diagnosis.

In this paper, we focus on estimating the bias of the GxE interaction when logistic regression is performed with the *observed* disease status as the dependent variable and the gene, environment, and their interaction as the independent variables. Our discussion builds on the literature that describes the bias of the main effects (i.e. gene or environment) in the presence of silent cases (Carroll et al, 2006) and, more specifically, Neuhaus’s (1999) approximation to the bias of the main effects when the data are collected using prospective sampling and analyzed in a prospective likelihood function.

Our paper proceeds as follows. First, in the Material and Methods section, we describe our notation and derive the theoretical approximation bias that results from ignoring the presence of silent disease. Next, in the Simulation Experiments section, we compare the theoretical approximation to empirical estimates of the bias across multiple scenarios. Then, we apply our approach to a Prostate Cancer GWAS (https://www.ncbi.nlm.nih.gov/projects/gap/cgi-bin/study.cgi?study_id=phs000207.v1.p1,Yeager et al, 2007). Finally, we conclude our paper with a brief Discussion section.

## MATERIALS AND METHODS

For individual *i*, let *G_i_* be the genotype, *X_i_* be the environmental variable potentially interacting with the genotype, and ⋅*Z_i_* be a vector of other environmental variables. Furthermore, let *D_i_* = {0,1} be a binary indicator of the *true*, and unobserved, disease status and let 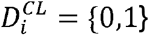 be a binary indicator of *clinically diagnosed* disease status. In the overall population, let *π*_0_ = pr(*D^CL^* = 0) and *π*_1_ = pr(*D^CL^* = 1),and in our study population let *n*_0_ be the number of controls (i.e. *D^CL^* = 0), *n*_1_ be the number of cases (i.e. *D^CL^* = 1), and n =n_0_ + n_1_. For clarity of presentation we suppose that all variables are binary, but the discussion could be easily extended to categorical variables, though the interpretation of GxE can then be notoriously difficult.

If *θ* is the frequency of minor allele *a* when the major allele is A, then the Hardy-Weinberg Equilibrium model (Hardy, 1908) states

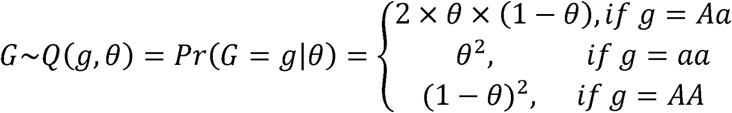

We assume that individuals with a clinical diagnosis have the true disease, i.e. pr(*D* = 1|*D^CL^* = 1),and that a substantial proportion of “controls” also have the true disease and that this proportion can vary by environmental factors: pr(*D* = 1|*D^CL^* = 0,*X*) = *τ*(*X*) > 0.

We next assume that the probability of the *true* disease follows a logistic model

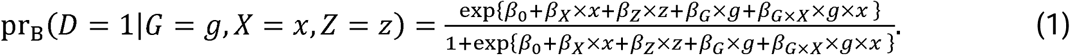

Define Β = (*β*_0_, *β_X_*, *β_Z_*,*β_G_*, *β_G_*_×*X*_) to be the vector of coefficients of interest.

The observed data are collected using retrospective sampling design, hence the likelihood function of the observed data is based on the probability *pr*[*G* = *g*, *X* = *x,Z* = *z*|*D^CL^* = *d^cl^*] and we define

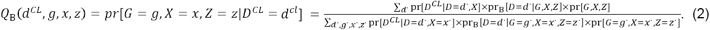

The usual analyses with the clinical diagnosis as an outcome variable and hence ignores presence of silent disease is based on the disease risk model

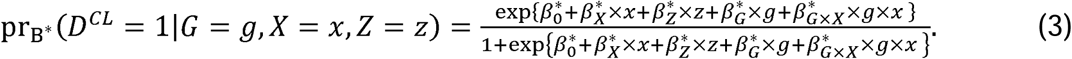

Estimation and inference in this setting is performed based on the likelihood function in the form

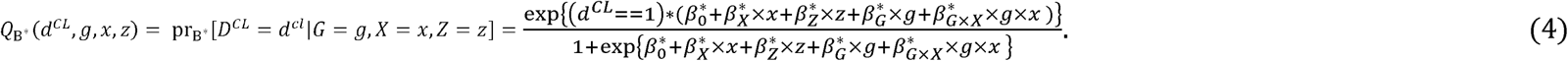

We are interested to find an analytic solution that relates parameters 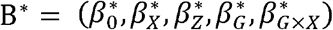 from the misspecified model (4) to the parameters Β = (*β*_0_, *β_X_*, *β_Z_*, *β_G_*, *β_G_*_×*X*_,from the *true* model (1)-(2).

The next steps are motivated by the developments in Kullback (1959), Neuhaus (1999). Kullback (1959) showed that parameters 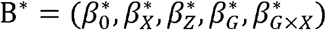 estimated in the misspesified model (4) converge to values that minimize the Kullback-Leibler divergence between the *true* and false models with expectations taken with respect to the *true* model, i.e.

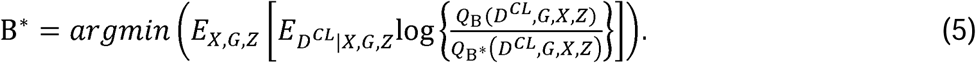

We define *γ*(*X*) = pr(*D^CL^* = 1|*D* = 1,*X*).

Derivations shown in Appendix arrive at the following approximation of the relationship between the parameters of the misspecified model (4) and the true model (1). For clarity of presentation we first assume that variable *Z* is not in the risk model. Generalization to include *Z* is described in Web-based supplementary materials.

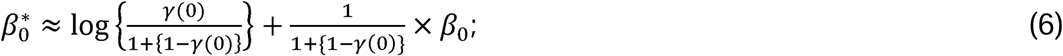

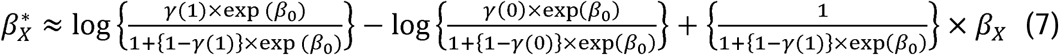

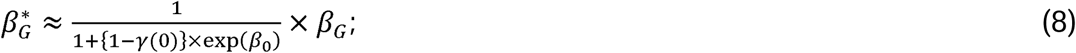

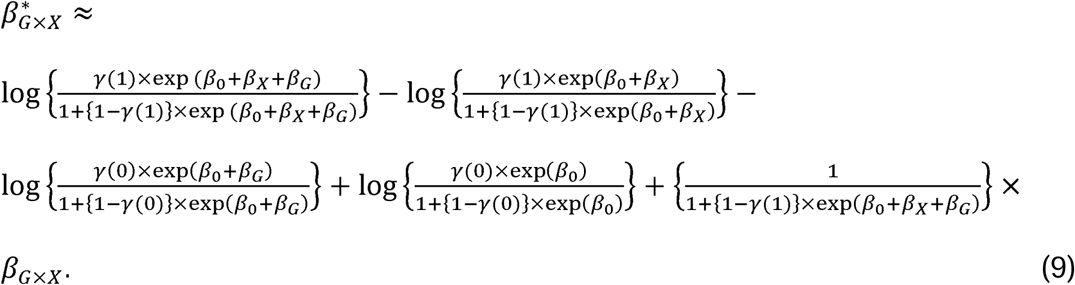

We now derive alternative formulation. In retrospective design cases and controls are sampled conditionally on the disease status. We therefore introduce an imaginary indicator of being selected into the study, ∆ = 1. Cases and controls are then selected into the study with probabilities 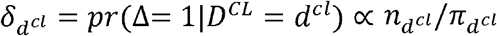. The true disease model then becomes

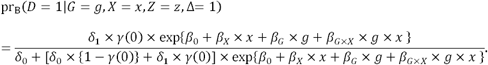

We then derive

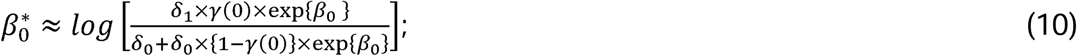

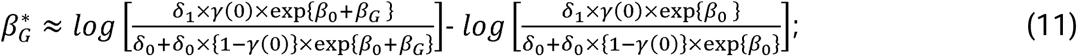

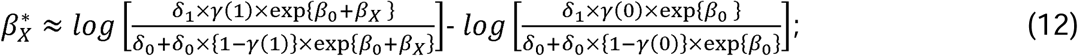

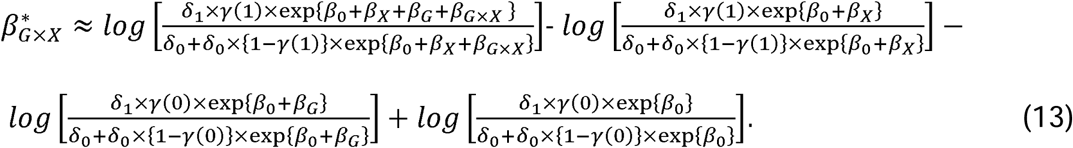

### Remarks

1. Appendix provides formulas (A11)-(A15) for the setting with environmental variable *Z* that does not interact with the SNP genotype and environmental variable *X*.
2. Appendix also provides formulas (A16)-(A21) for the setting when the environmental variable *Z* interacts with the environmental variable *X*, but does not interact with the SNP genotype.
3. When the clinical diagnosis and pathologic disease status correspond, i.e.*γ* (0) = *γ*(1) = 1, then all parameter estimates are unbiased.
4. If *β_G_* = 0, then 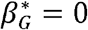. Hence the usual logistic regression yields a consistent estimate of the null *β_G_*.
5. If *β*_0_ = 0, then 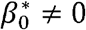. Similarly, if *β_X_* = 0, then 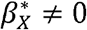; and if *β_G_*_×*X*_ = 0, then 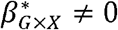. Hence the usual logistic regression does not yield a consistent estimate of the null effect *β*_0_, *β_X_*, *β_G_*_×*X*_.
6. If *β_G_* = 0 and *β_X_*_×*G*_ = 0 then 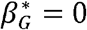 and 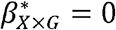. Hence the usual logistic regression yields consistent estimate of the null *β_G_* and *β_X_*_×*G*_.
7. If the misclassification is non-differential, i.e. *γ*(0) = *γ*(1); and if *β_X_* = 0, then 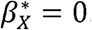. That is then the usual logistic regression model yields consistent estimate of the null effect *β_X_*.
8. If the misclassification is non-differential, i.e. *γ*(0) = *γ*(1); *β_X_* = 0, *β*_G_ = 0, *β_X_*_×*G*_ = 0 simply 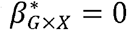. That is then the usual logistic regression model yields consistent estimate of the null effect of *β_G_*_×*X*_.
9. Taylor series expansion of (10)-(13) around the true parameters equal to zero arrives to (6)-(9).

## SIMULATION EXPERIMENTS

We first perform a set of simulation studies to investigate a false positive rate for *β_G_* _×*X*_ estimates. We define the false positive rate to be the proportion of p-values ≤0.05 from the usual logistic regression with the clinical diagnosis as an outcome variable across 10,000 studies. We simulate *X* to be binary with frequency 0.488 and *G* with frequency 0.10. Next, we simulate the *true* disease status according to the risk model with coefficients *β*_0_ = 1,1, *β_G_* = log(1), log(1.5), log(2), log(2.5), log(3), log(3.5), log(4), log(4.5), *β_X_* = log (2), *β_G_*_×*X*_ = 0. To simulate the clinical diagnosis we define the clinical-analyses, i.e. pr (*D* =1|*D^CL^* = 0,X) = 0.252 and 0.389 for *X* = 0,1. We simulate pathological diagnoses relationship to be as in the Prostate Cancer data datasets with *n*_0_ = *n*_1_ = 3,000, *n*_0_ = *n*_1_ = 1,000. False positive rates shown in **Table 1** indicate that the rate is the nominal when main effect of the genotype is zero, and increases as the value of *β_G_* increases. When frequency of the *true* disease is higher (*β*_0_ _1 *vs*. –1) then overall the false positive rates are lower. For example, in a study with *n*_0_ = *n*_1_ = 3,000, when *β_G_* = log(3) = 1.1, the rates are 0.19 and 0.14, when *β*_0_ = – 1and 1, respectively. The false discovery rates are persistently elevated in studies with *n*_0_ = *n*_1_ = 10,000.

**Table 1.** False positive rate for *β_G_*_×*X*_ estimates. We define the false positive rate to be the proportion of p-values ≤0.05 from the usual logistic regression with the clinical diagnosis as an outcome variable across 10,000 studies. We simulate *X* to be binary with frequency 0.488 and *G* with frequency 0.10. Next, we simulate the *true* disease status according to the risk model with coefficients *β*_0_ = –1,1, *β*_G_ = log(1), log(1.5), log(2), log(2.5), log(3), log(3.5), log(4), log (4.5), *β_X_* = log(2), *β_G_*_×*X*_ = 0. To simulate the clinical diagnosis we define the clinical-pathological diagnoses relationship to be as in the Prostate Cancer data analyses, i.e. pr(*D* = 1|*D^CL^* = 0, X) = 0.252 and 0.389 for *X* = 0, 1. We simulate datasets with *n*_0_ = *n*_1_ = 3,000, *n*_0_ = *n*_1_ = 1,000.

| $\beta_G =$ | | Log(1)<br>=0 | Log(1.5)<br>=0.41 | Log(2)<br>=0.69 | Log(2.5)<br>=0.69 | Log(3)<br>=1.1 | Log(3.5)<br>=1.3 | Log(4)<br>=1.4 | Log(4.5)<br>=1.5 |
| --- | --- | --- | --- | --- | --- | --- | --- | --- | --- |
| $n_0 = n_1$<br>= 3,000 | $\beta_0 = -1$ | 0.047 | 0.066 | 0.098 | 0.14 | 0.19 | 0.24 | 0.27 | 0.32 |
| | $\beta_0 = 1$ | 0.05 | 0.057 | 0.087 | 0.11 | 0.14 | 0.16 | 0.19 | 0.22 |
| $n_0 = n_1$<br>= 10,000 | $\beta_0 = -1$ | 0.051 | 0.11 | 0.23 | 0.31 | 0.41 | 0.53 | 0.68 | 0.72 |
| | $\beta_0 = 1$ | 0.05 | 0.08 | 0.18 | 0.26 | 0.36 | 0.44 | 0.51 | 0.57 |

We conducted simulation studies to evaluate the accuracy of the theoretical approximation that we derived in (6)-(9) and in the Appendix. These studies are presented in Web-based Supplementary Materials.

We next describe the magnitude of bias in estimates of *β_G_*_×*X*_ = 0 for various frequencies of the clinical diagnosis and the *true* disease state in the population. We simulate *X* as Bernoulli with frequency 0.488 and *G* as Bernoulli with frequency 0.10. We next simulate the *true* disease status using coefficients *β*_0_ = – 3.5, –3, –2.5, –2, –1.5, –1, –0.5,0,0.5,1,1.5,2,2.5,3,3.5; *β_G_* = log(1.5) = 0.41, *β_X_* = log(3) = 1.099, *β_G_*_×*X*_ = 0. We next simulate the clinical diagnosis with frequencies

*γ*(0) = pr (*D^CL^* = 1|*D* = 1, *X* = 0) = 0.000001,0.0001,0.001,0.005,0.01,0.10, and *γ*(1) = pr(*D^CL^* = 1|*D* = 1, *X* = 1) = 0.000001,0.0001,0.001,0.005,0.01,0.10. We then estimate bias in estimates of *β_G_*_×*X*_ using (13) for each of the above settings.

Shown on **Figure 2** are frequencies of the *true* probability of disease across values of *β*_0_ on the x-axis. **Figure 3** presents probabilities of the clinical diagnosis across values of *β*_0_ on the x-axis, values of *γ*(1) on the panels, and values of *γ*(0) indicated by color. We note that the setting of prostate cancer example corresponds to the values of *β*_0_ around −2 and *γ*(0) ≈ 1,.0.000001. Bias in the estimates of *β*_***G***×*X*_ shown on **Figure 4** differs across values of *β*_0_, *γ*(0), *γ*(1). Magnitude of bias can be substantial and is usually smaller when *γ*(0) = *γ*(1).

**Figure 2:**
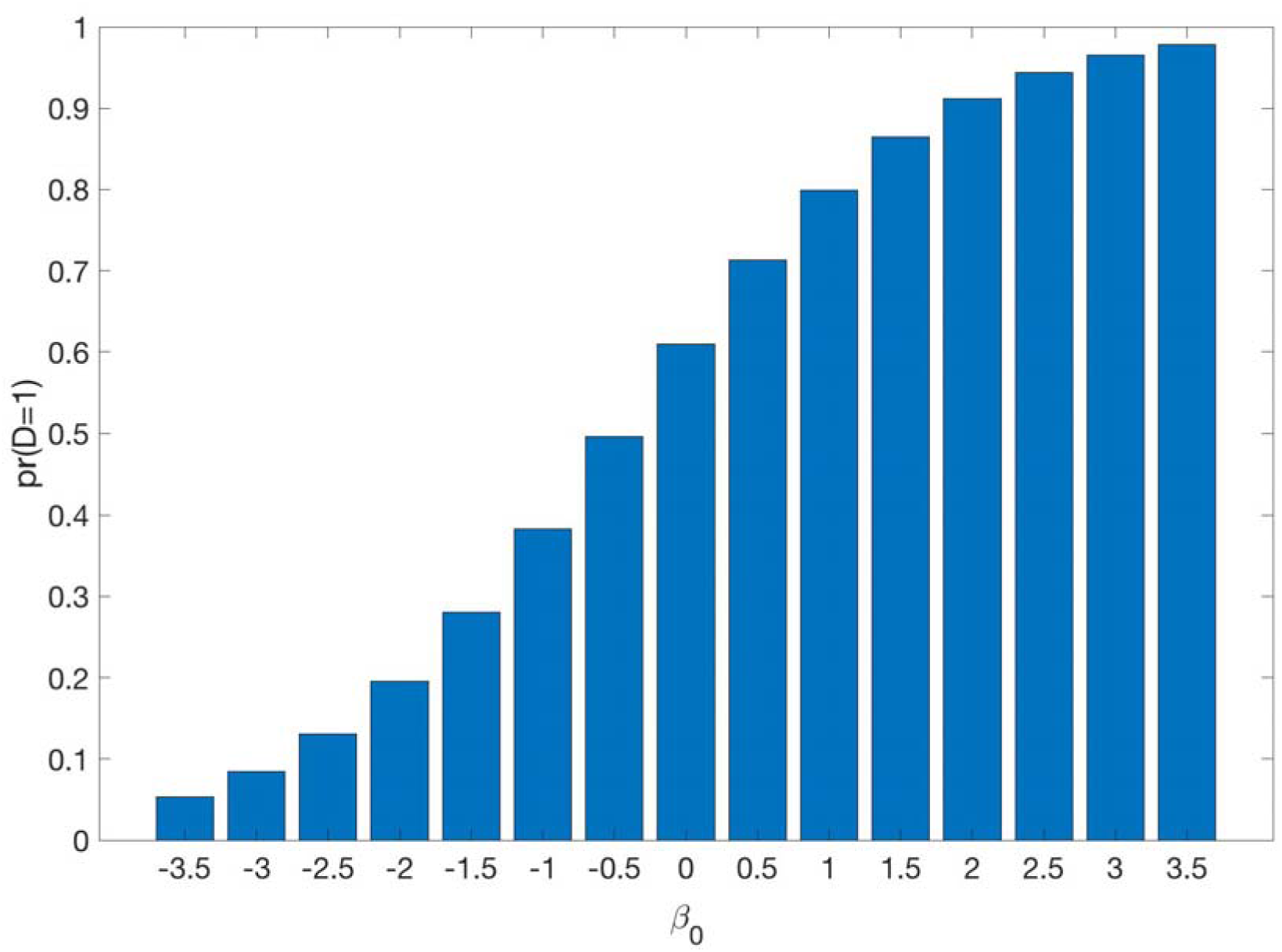
Frequencies of the true disease status in the population for various values of the intercept. We simulate *X* as Bernoulli with frequency 0.488 and c as Bernoulli with frequency 0.10. We next simulate the *true* disease status using coefficients *β*_0_ = –3.5, –3, –2.5, –2, –1.5, –1, –0.5, 0, 0.5, 1, 1.5, 2, 2.5, 3, 3.5; *β_G_* = logC1.5) = 0.41, *β_X_* = log(3) = 1.099, *β_G_*_×*X*_ = 0.

**Figure 3:**
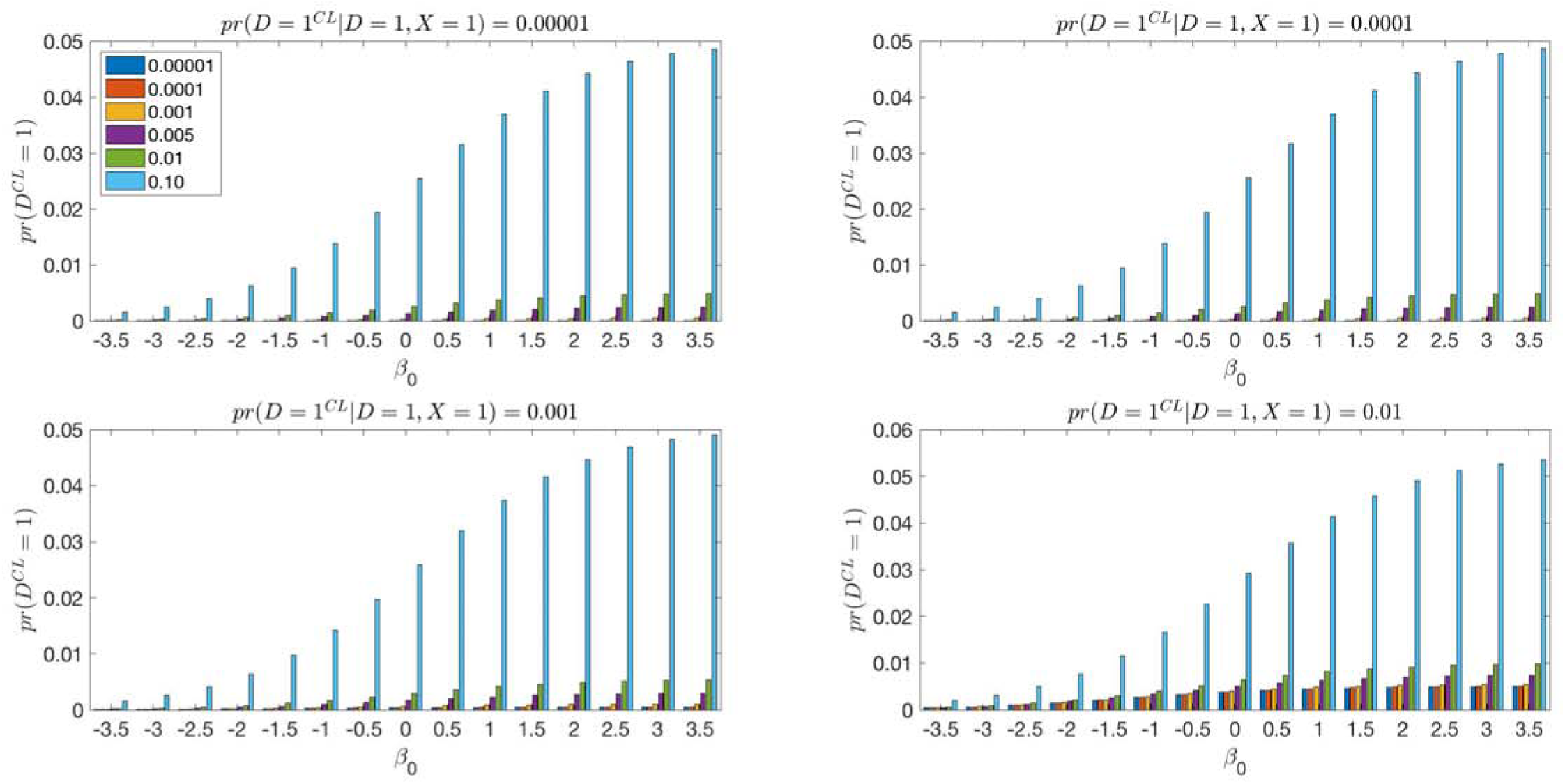
Frequencies of the clinical diagnosis in the population for various values of the intercept along the x-axis, values of *γ*(1) = pr(*D^CL^* = 1|*D* = 1, *X* = 1) across the panels and values of *γ*(1) = pr(*D^CL^* = 1|*D* = 1, *X* = 0) as indicated by color. We simulate *X* as Bernoulli with frequency 0.488 and *G* as Bernoulli with frequency 0.10. We next simulate the *true* disease status using coefficients *β*_0_ = –3.5, –3, –2.5, –2, –1.5, –1, –0.5, 0, 0.5, 1, 1.5, 2, 2.5, 3, 3.5; *β_G_* = log(1.5) = 0.41, *β_X_* = log(3) = 1.099, *β_G_*_×*X*_ = 0. We next simulate the clinical diagnosis with frequencies *γ*(0) = pr(*D^CL^* = 1|*D* = 1, *X* = 0) = 0.000001, 0.0001, 0.001, 0.005, 0.01, 0.10, and *γ*(1) = pr(*D^CL^* = 1|*D* = 1, *X* = 1) = 0.000001, 0.0001, 0.001, 0.005, 0.01, 0.10.

**Figure 4:**
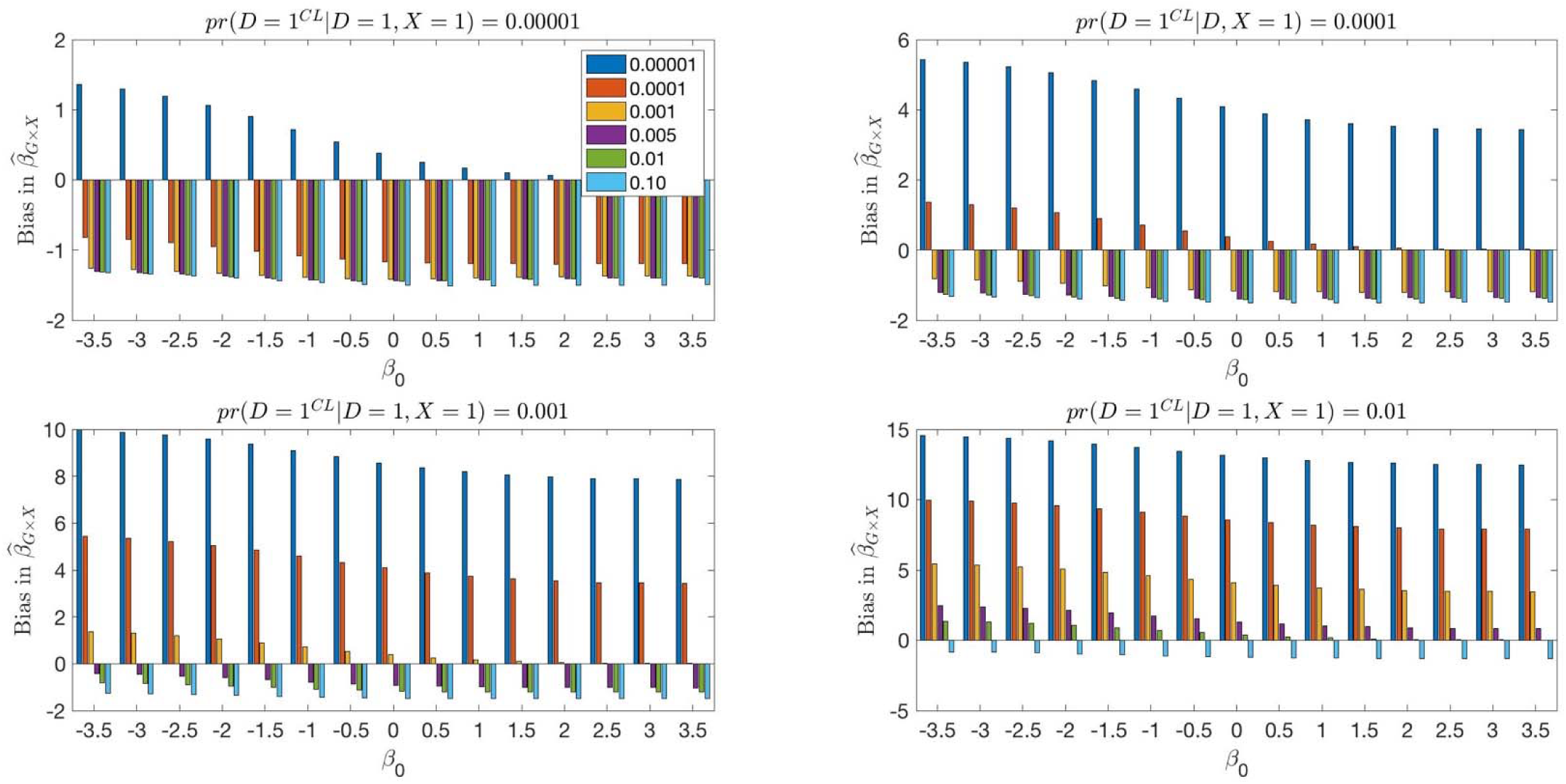
Bias in the estimates of *β_G_*_×*X*_ for various values of the intercept along the x-axis, values of *γ*(1) = pr(*D^CL^* = 1|*D* = 1, *X* = 1) across the panels and values of *γ*(1) = pr(*D^CL^* = 1|*D* = 1, *X* = 0) as indicated by color. We simulate *X* as Bernoulli with frequency 0.488 and *G* as Bernoulli with frequency 0.10. We next simulate the *true* disease status using coefficients *β* = –3.5, –3, –2.5, –2, –1.5, –1, –0.5, 0, 0.5, 1, 1.5, 2, 2.5, 3, 3.5; *β_G_* = log(1.5) = 0.41, *β_X_* = log(3) = 1.099, *β_G_*_×*X*_ = 0. We next simulate the clinical diagnosis with frequencies *γ*(0) = pr(*D^CL^* = 1|*D* = 1, *X* = 0) = 0.000001, 0.0001, 0.001, 0.005, 0.01, 0.10, and *γ*(1) = pr(*D^CL^* = 1|*D* = 1, *X* = 1) = 0.000001, 0.0001, 0.001, 0.005, 0.01, 0.10.

## PROSTATE CANCER DATA ANALYSES

We performed GxE analyses for Prostate Cancer using data collected as part of the Prostate, Lung, Colon and Ovarian (PLCO) Screening trial (dbGAP: https://www.ncbi.nlm.nih.gov/projects/gap/cgi-bin/study.cgi?study_id=phs000207.v1.p1, study accession phs000207.v1.p1,Yeager et al, 2007). The study included 965 cases and 1,035 controls of European ancestry with 550,000 genotyped SNPs. The number of cases in 50-69 and 70+ year age groups were 636, 329, respectively; the number of controls in the same groups were 727 and 308. Furthermore, 11.3% of cases and 6.2% of controls had a family history of prostate cancer. In the following analyses, we focus on SNPs serving mitochondria. We mapped the SNPs onto human chromosomes using NCBI dbSNP database https://www.ncbi.nlm.nih.gov/projects/SNP/ and recorded chromosome location, proximal gene or genes in the gene structure location (e.g. intron, exon, intergenic, UTR). Based on these data, we inferred 1,867 SNPs serving mitochondria according to MitoCarta database (https://www.broadinstitute.org/scientific-community/science/programs/metabolic-disease-program/publications/mitocarta/mitocarta-in-0).

For each of the 1,867 SNPs, we assumed the relationship between the *true* disease status and the combination of SNP, family history, and age can be described by logistic regression, i.e. model (3).

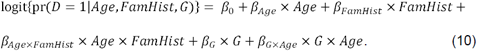

We assumed the relationship between clinical disease status and the *true* disease status is pr(*D* = 1|*D^C^* = 0,Age) = 0.252 and 0.389 for age groups of 50-69 and 70+, respectively (Jahn et al, 2015). We suppose that the clinical diagnosis is correct for all cases (Canto and Slawin, 2002).

We first estimate the coefficients using the usual logistic regression model without considering the correction for the silent disease. Then we estimate the corresponding coefficient of the *true* model from the approximation derived in Appendix (A16)-(A21) with the consideration of the relationship between the clinical disease status and the *true* disease status.

The usual logistic regression estimate for the intercept is −0.19, while the approximation to the bias is −0.60. In the usual logistic regression *β*̂_*FamHist*_ = 0.60 and the usual and the bias is estimated to be −0.23. Across all SNPs, the usual estimate of estimate of *β_Age_* is on average 0.21, while the bias is approximated to be −0.68; and the usual estimate of *β_Age_*_×*FamHist*_ is on average 0.08, while the bias is approximated to be −0.82. Shown on **Figure 5A** is the histogram of bias in *β_G_* across 1,975 SNPs that ranges from −0.19 to 0.20 with an average of 0.0042. Shown on **Figure 5B** is the histogram of bias in *β_G_*_×*Age*_ ranging from −1.87 to 0.81 with an average of −0.07.

**Figure 5.**
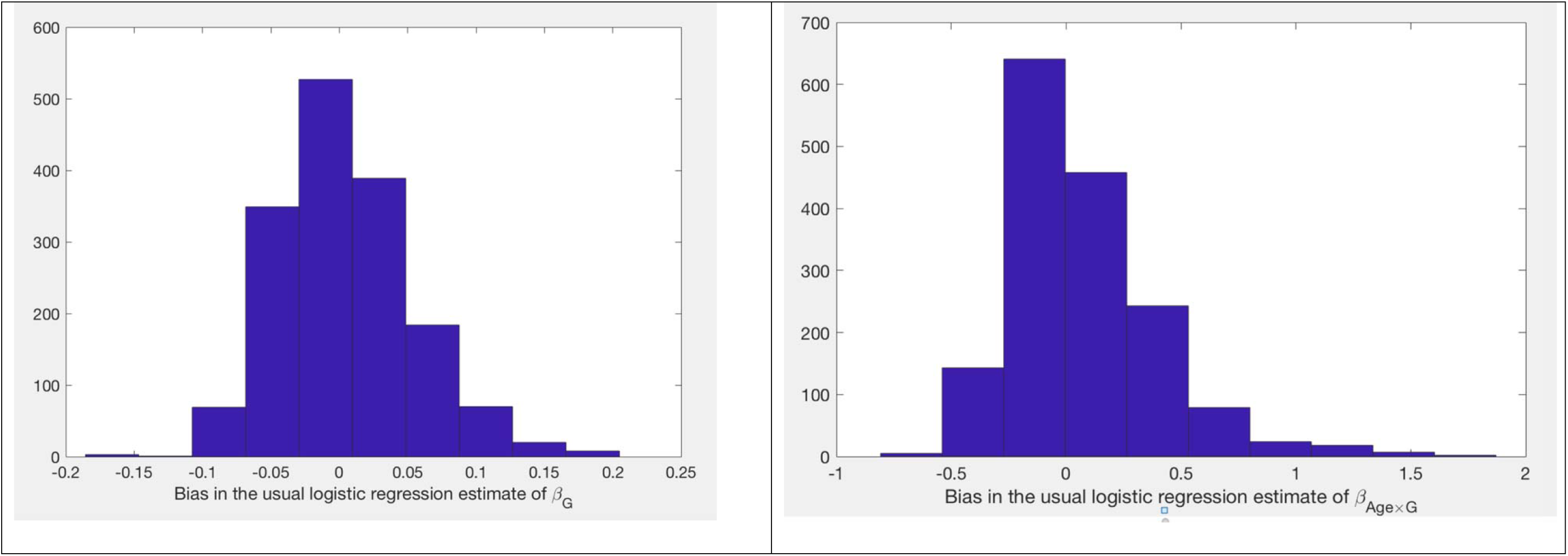
A (left panel): Histogram of the bias of the usual logistic regression estimate of in Prostate Cancer dataset. The bias is approximated using equations (A16)-(A21) and **Figure 5B (right panel):** Histogram of the bias of the usual logistic regression estimate of in Prostate Cancer dataset. The bias is approximated using equations (A16)-(A21).

## DISCUSSION

We derived a general and convenient theoretical approximation to the bias in GxE parameter estimates for studies where a substantial fraction of the controls are undiagnosed cases. In case-control studies the usual logistic regression model produces biased estimates either because the presence of the latent cases is ignored, or because the sampling design is misspecified (analysis of case-control data by a prospective likelihood function while the data was collected retrospectively), or both.

While we have recently proposed a solution that eliminates the bias (Lobach et al, 2018), the implementation requires optimization of a complex non-linear equation. The approximation that we’ve developed provides convenient estimates of the bias and a clear explanation of how all parameter estimates can be biased. The presence of the silent disease distorts the true link between the GxE interaction and the true disease status.

In the analyses of Prostate Cancer, we note that bias in GxE estimates can be in either direction resulting in either under- or over-estimation of the magnitude of the effect. Similarly, the bias in *β_G_* manifested itself in either direction.

The approximation that we’ve developed is a first order Taylor series expansion of a solution that minimizes Kullback-Leibler divergence criteria between the true and the misspecified models. While the Kullback-Leibler divergence could have multiple local minima, in the extensive simulations studies that we considered the numerical optimization did find the minimum that was accurate relative to the empirical estimates. The theoretical approximation can be improved by deriving further order Taylor series expansions.

We note that the bias in GxE generally decreases as the frequency of the true disease and the clinical diagnosis decrease. The magnitude of bias, however, can be substantial even when the disease is common, similarly to what has been described for common diseases in trio designs (Peyrot et al, 2016). Specifically, when frequency of the silent disease varies by the environmental variable. The bias is more elastic as a function of how frequencies of the environmental variable are different by the environment, i.e. there is more change in the parameter estimates.

The proposed analyses rely on knowing the estimates of silent disease in the population subgroups. These estimates are often available in epidemiologic studies or can be estimated in an internal reliability study. For example, in the study of Prostate Cancer, the rates of silent disease are estimated based on a sample of size 3,799 US Whites and Europeans. If the estimates of the rates are with high uncertainty, the approximation that we derived provides a convenient and general formulae to understand how much the estimates can change across various settings defined by frequencies of the silent disease and frequencies of the disease and the clinical diagnoses in the population. If the proportion of silent cases is not known, the approximations that we derived provide a simple way to examine potential bias across various rates for silent disease that are plausible. Such analyses might inform how elastic the effect estimates can be for a given value of the estimate and frequency of the clinical diagnosis.

The goal for exploring GxE is to investigate if the effect of a genetic variables varies by non-genetic (environmental) variables. We described one source of bias in estimates of GxE, namely due to ignoring presence of silent cases. Other biases in the estimates have been noted in literature. For example, Keller (2014)note the widespread bias in GxE due to inappropriately controlling for covariates while studying GxE. We have recently analyzed bias in the estimates due to omitting GxE (Lobach, 2018).

## Acknowledgements

Prostate cancer dataset was downloaded from the database of genotypes and phenotypes (https://dbgap.ncbi.nlm.nih.gov), study accession number phs000297.v1.p1.

https://www.ncbi.nlm.nih.gov/projects/gap/cgi-bin/study.cgi?study_id=phs000207.v1.p1.

We thank Ivan Belousov for help with the computations.

